# Distinct features of human myeloid cell cytokine response profiles identify neutrophil activation by cytokines as a prognostic feature during tuberculosis and cancer1

**DOI:** 10.1101/775296

**Authors:** Joseph C. Devlin, Erin E. Zwack, Mei San Tang, Zhi Li, David Fenyo, Victor J. Torres, Kelly V. Ruggles, P’ng Loke

**Affiliations:** Sackler Institute, Department of Medicine, New York University School of Medicine, New York, NY, USA; Department of Microbiology, New York University School of Medicine, New York, NY 10016, USA; Division of Translational Medicine, Department of Medicine and Department of Microbiology, New York University School of Medicine, New York, NY, USA; Applied Bioinformatics Laboratories, New York University School of Medicine, New York, NY, USA; Department for Biochemistry and Molecular Pharmacology, NYU Langone Health, New York, NY 10016, USA; Institute for Systems Genetics, NYU Langone Health, New York, NY 10016, USA

## Abstract

Myeloid cells are a vital component of innate immunity and comprise of monocytes, macrophages, dendritic cells and granulocytes. How myeloid cell lineage affects activation states in response to cytokines remains poorly understood. The cytokine environment and cellular infiltrate during an inflammatory response may contain prognostic features that could predict disease outcome. Here we analyzed the transcriptional responses of human monocytes, macrophages, dendritic cells and neutrophils in response to stimulation by IFN-γ, IFN-β IFN-λ, IL-4, IL-13 and IL-10 cytokines, to better understand the heterogeneity of activation states in inflammatory conditions. This generated a myeloid cell cytokine specific response matrix that can infer representation of myeloid cells and the cytokine environment they encounter during infection and in tumors. Neutrophils were highly responsive to type 1 and type 2 cytokine stimulation but did not respond to IL-10. We identified transcripts specific to IFN-β stimulation, whereas other IFN signature genes were upregulated by both IFN-γ and IFN-β. When we used our matrix to deconvolute blood profiles from tuberculosis patients, the IFN-β specific neutrophil signature was reduced in TB patients with active disease whereas the shared response to IFN-γ and IFN-β in neutrophils was increased. When applied to glioma patients, transcripts of neutrophils exposed to IL-4 or IL-13 and monocyte responses to IFN-γ or IFN-β emerged as opposing predictors of patient survival. Hence, by dissecting how different myeloid cells respond to cytokine activation, we can delineate biological roles for myeloid cells in different cytokine environments during disease processes, especially during infection and tumor progression.

## INTRODUCTION

While there has been rapid recent progress in understanding the ontogeny of myeloid cells, including monocytes, macrophages, dendritic cells and granulocytes in recent years, the heterogeneity of activation states between these different cell types remains poorly understood. Single cell RNA seq technologies of inflamed tissues has begun to provide an appreciation for the heterogeneity of activation states for different myeloid cells, however these cells typically encounter a complex mixture of cytokines in their tissue microenvironment. The overall status of immune cells in a particular tissue or in blood circulation in disease conditions is an important indicator of disease state. Transcriptional profiles of immune cells have thus been used to define gene expression signatures that could potentially guide personalized clinical decision-making through patient stratification and evaluation of disease-associated gene expression changes. However, in most cases, transcriptional profiles are generated from bulk tissues or whole blood, masking changes in the transcriptomic composition of specific cell types. Recently, computational approaches have been developed to infer leukocyte compositions in bulk tissue transcriptomes based on cell-type specific reference gene expression signatures (1). One such study found that the ratio of tumor-associated neutrophils and plasma cell signatures was predictive of survival for various solid tumors (2). While this strategy enables the deconvolution of immune cell types infiltrating different tissues, the environmental conditions they encounter as they infiltrate the tissues is not yet known.

Identifying specific transcriptional programs in myeloid cells may facilitate the discovery of biomarkers and targets for therapies for a variety of diseases. Both granulocytic myeloid cells (e.g. neutrophils, eosinophils and basophils) and monocytic myeloid cells are important innate immune components of the inflammatory infiltrate, being almost universally present in any disease condition. They are all critical not just for protection against pathogens but also for tissue remodeling and maintenance of tissue homeostasis. The same differentiation processes that guide the physiologically necessary function of these cells are also responsible for the pathological accumulation of these cells under certain inflammatory conditions. For example, myeloid derived suppressor cells (MDSCs) can play pathological roles in cancer, as well as other inflammatory settings where they accumulate and differentiate (3).

The cytokine environment is a critical determinant of immune cell activation phenotypes and the response of diverse immune cells to the different cytokines is not well understood. Further, cell types respond differentially to various cytokine stimulation conditions to express distinct transcriptional signatures. This may be due to differences in chromatin state and cytokine receptor expression levels that determine, for example, how macrophages and dendritic cells respond to IL-10 stimulation as compared to IFN-γ stimulation (4, 5). While there have been experimental studies whereby transcriptional response has been assessed in specific immune cell types following exposure to assorted cytokines, we are not aware of a systematic comparison of diverse myeloid cell types in response to a wide variety of different cytokine stimulation conditions. Here, we compare the transcriptional response of primary human macrophages, dendritic cells, monocytes and neutrophils to stimulation with a cytokine panel consisting of IL-4, IL-10, IL-13, IFN-γ, IFN-β, and IFN-λ. These signatures were then used to infer the signature of specific immune cell types responding to specific cytokine environments from bulk transcriptomic data. This method allows us to infer not only the type of immune cells present in a bulk tissue or blood but also the cytokine environment which they are likely encountering. We have successfully identified 12 myeloid cell-cytokine stimulation signatures and correlated both *Mycobacterium tuberculosis* infection status and glioma cancer outcome with these specific signatures.

## MATERIALS AND METHODS

### Cell Isolation and Differentiation Protocol

Primary human polymorphonuclear neutrophils (PMNs) and peripheral blood mononuclear cells (PBMCs) from anonymous, healthy donors (New York Blood Center) were isolated by Ficoll gradient separation as previously described (6). CD14+ monocytes were then isolated from the PBMC fraction by positive selection. In brief: PBMCs were resuspended in MACS buffer (PBS + 0.05% BSA + 2 mM EDTA) at a concentration of 1×10_8_ PBMCs per 950 μL. 50 μL of CD14+ microbeads (Miltenyi Biotec) were added for every 1×10_8_ PBMCs. Cells were incubated for 20 minutes at 4°C, washed, and filtered through a cell strainer. The cells were run on an AutoMACS Pro (Miltenyi Biotec) using the ‘Posselds’ program. Monocytes were used directly after sorting. Monocyte-derived dendritic cells (DCs), and Monocyte-derived macrophages were differentiated from CD14+ monocytes by culturing the cells for 4 days at 37°C and 5% CO_2_ in RPMI medium supplemented with 10% FBS, 10 mM HEPES, 100 U/mL penicillin, 100 μg/mL streptomycin with either 110 U/mL granulocyte-macrophage colony-stimulating factor (GM-CSF) (Leukine; Sanofi) and 282 U/mL interleukin-4 (IL-4) (Affymetrix, eBioscience) for DCs or 280 U/mL GM-CSF for macrophages. Media was replenished with fresh cytokine on day 2.

### Cell Stimulation Protocol

Differentiated cells were resuspended in clear RPMI + 10% FBS. 1×10_5_ cells were added to each stimulation well. Stimulations comprised of buffer control (PBS + 0.01% Glycerol), 500 U/mL IFN-β1a (Carrier Free; R&D Systems), 10 ng/mL IFN-γ (Carrier Free, R&D Systems), IFN-λ2 (Carrier Free, R&D Systems), 1000 IU/mL IL-4 (Carrier Free, Life Technologies), 100 IU/mL IL-10 (Carrier Free, Life Technologies), and 100 IU/mL IL-13 (Carrier Free, R&D Systems). Plates were spun for 5 minutes at 1200 rpm and incubated for 4 hours at 37°C and 5% CO_2_. Cells were then washed with PBS. Cells were resuspended in RLT buffer (Qiagen) and vortexed for 1 minute before being placed at −80°C. RNA for each donor was then isolated with the RNeasy Plus Mini Kit (Qiagen) following the protocol with on column DNAse Digestion (Qiagen).

### Gene Expression Analysis

Libraries were generated for each donor using the CelSeq2 protocol (7) and were sequenced on Illumina Hi-Seq. Reads were mapped by Bowtie2.3.1 (8) to the hg38 reference genome and uniquely mapped indices (UMI) determined by HTSeq-counts (9). Differential expression analysis was performed in R (v3.5.1) using DESeq2 (10). Compared to buffer controls, differentially expressed genes were considered significant with Log2 fold change greater than 2 and adjusted p-value less than 0.05.

### Self-Organizing Map and Outlier Analysis

Self-organizing map (SOM) analysis (11) was performed on the list of 571 differentially expressed genes using the R statistical programming language. SOM analysis was performed individually for each cell type with the R package Kohonen (12) at default parameters. According to 16 identified SOM clusters outlier analysis was performed to identify specific gene expression patterns. A gene was considered an outlier with an expression level 1.5 times greater than the median expression level across all conditions in at least two out of the three donors (13). 131 of 571 genes were found to meet these criteria in 12 of the possible 16 cell type and stimulation conditions.

### Cell type deconvolution through CIBERSORT

Source code for the CIBERSORT deconvolution algorithm, https://cibersort.stanford.edu/, was obtained from the developers and implemented in the R statistical programming language (14) All input bulk datasets were obtained as normalized count tables when available. If not normalized datasets were scaled and quantile normalized according to the default CIBERSORT functions. Our MCCS basis matrix was supplied as the average normalized expression level across the three donors for our 131-gene set. The basis matrices for immunoStates (15) and LM22 (1) were obtained from the respective publications. CIBERSORT was run according to default parameters in all cases with 100 permutations.

### *M. tuberculosis* sample collection and normalization

Conducting a literature search for all available TB infection studies with publicly available data yielded 8 microarray and 5 RNA-Seq studies, with the following accession numbers; GSE19491, GSE28623, GSE37250, GSE39939, GSE39940, GSE40553, GSE41055, GSE56153, GSE101705, GSE107995, GSE79362, GSE89403, GSE94438 (16–28). See **Table S2** for full sample details. Microarray studies were obtained as scaled expression values as downloaded from GEO. RNA-Seq studies were obtained as edgeR (29) normalized count tables.

### LASSO modeling and feature selection for patient survival in primary gliomas

RSEM normalized count tables for all primary glioma samples available in the TCGA database were obtained through the TCGA2STAT R package (30). Additional sample metadata was also obtained from Ceccarelli et al. (31). Samples were randomly split into a training set and a test set with an 80/20 split depending on the vital status at the 2-year or 5-year model. Additionally, survival status was balanced as much as possible between the test and train sets to improve model predictions. In the 2-year model there were 264 samples (133 alive, 131 deceased) in the training set and 66 (32 alive, 34 deceased) samples in the test set. And in the 5-year model there were 358 samples (168 alive, 190 deceased) in the training set and 90 (56 alive, 34 deceased) samples in the test set. Prior to modeling the samples were scaled with min-max normalization by normalizing the gene expression levels for each sample between 0 and 1. The sample breakdowns were subject to a logistic least absolute shrinkage and selection operator (LASSO) model with 7-fold cross validation repeated 10 times using the R package caret (32). Area under the receiver operator curve (AUC) and precision recall curves were used to assess model performance by the default functions in caret (32). Additionally, feature importance was assessed by the caret importance function, varImp, which measures the regression coefficients for each gene supplied to the model.

### Availability of data and material

Gene expression data is deposited in GEO under the accession number GSE131990. The TB infection studies of publicly available data includes 8 microarray and 5 RNA-Seq studies, with the following GEO accession numbers; GSE19491, GSE28623, GSE37250, GSE39939, GSE39940, GSE40553, GSE41055, GSE56153, GSE101705, GSE107995, GSE79362, GSE89403, GSE94438 (16–28). RSEM normalized count tables for all primary glioma samples are available in the TCGA database (https://portal.gdc.cancer.gov/) and were obtained through the TCGA2STAT R package (30). Additional sample metadata was also obtained from Ceccarelli et al. (31).

## RESULTS

### Myeloid cells respond to cytokine stimulation with cell type specific transcriptional profiles

In order to better understand how different human myeloid cells respond to activation by different types of cytokines, we set out to compare the transcriptional profiles attained through RNA-Seq of monocytes, neutrophils, macrophages and dendritic cells from the same healthy donors in response to stimulation by type 1 cytokines (IFN-γ, IFN-β and IFN-λ), type 2 cytokines (IL-4 and IL-13) and the regulatory cytokine IL-10. Neutrophils and monocytes were stimulated directly after isolation from blood leukopaks whereas macrophages and dendritic cells were stimulated after a 4-day differentiation period from the isolated monocytes (**Fig. 1A**). RNA was isolated 4 hours after stimulation for each of the four different cell types and stimulation conditions including an unstimulated buffer control for each cell type. Donor to donor differences had a much smaller effect on transcriptional profiles than differences between cell types (**Fig. S1**). We next identified genes that were significantly upregulated in individual cytokine stimulations relative to the unstimulated condition for each cell type. For example, with macrophages, we identified a total set of 341 genes that were significantly upregulated, log2 fold change greater than 2 and FDR less than 0.05, by at least one cytokine relative to the unstimulated control samples. Monocytes upregulated 197 genes; dendritic cells upregulated 199 genes; and neutrophils were highly responsive and upregulated 274 genes in response to cytokine stimulation (**Fig. 1C**). We then combined all of these lists for a total of 571 genes that are upregulated by at least one cytokine in at least one myeloid cell type. Principle component analysis (PCA) based on these genes indicated that each cell type engages a distinct transcriptional programming for each cytokine stimulation (**Fig. 1B**). 35% of the explained variation along the first principle component was strongly associated with cell type identity. Within each myeloid cell type, it is clear that type 2 cytokines IL-4 and IL-13 triggered shared transcriptional programs, whereas the type 1 cytokines IFN-β and IFN-γ triggered a similar set of upregulated genes (**Fig. 1C**). An IL-10 induced signature was observed in macrophages, dendritic cells and monocytes but completely absent in neutrophils. Interestingly, neutrophils had a robust response to other cytokines including a small subset of genes induced by IFN-λ, which was not observed in the other cell types (**Fig. 1C**).

**Figure 1.**
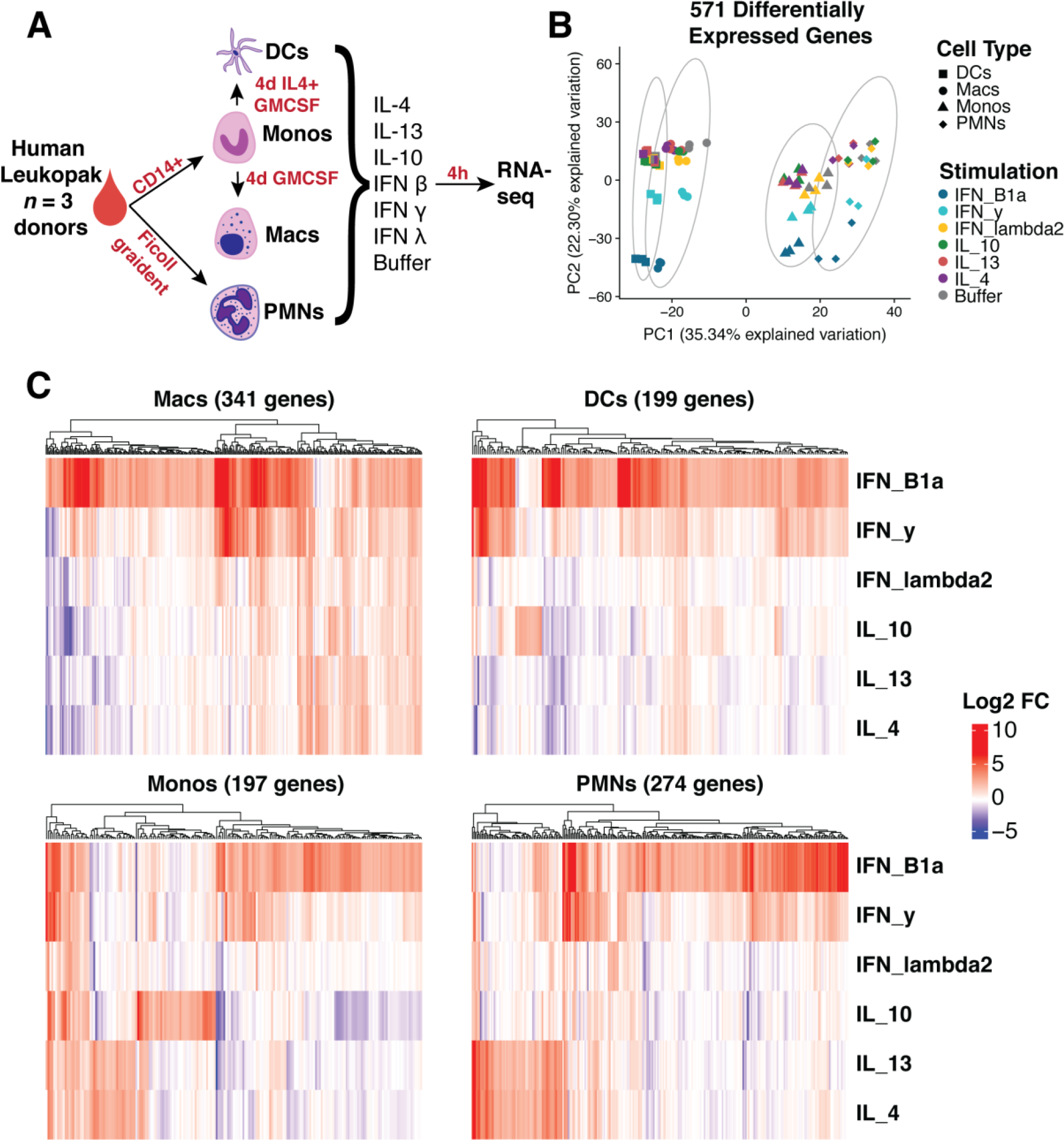
Transcriptional profiling indicates myeloid cell lineages respond strongly to cytokine stimulation. (A) Schematic of experimental workflow. Four different lineages of myeloid cells were isolated (PMNs and Monocytes) and differentiated (Macrophages and Dendritic Cells) from the same leukopaks from 3 healthy human donors. The cells were stimulated with a panel of six cytokines, as listed, and profiled for gene expression. (B) Principle component analysis of 571 genes determined by differential expression analysis compared to buffer condition. (C) Heatmaps of log2 fold change of differentially expressed genes in each cell type. Genes were considered significant with Log2 fold change greater than 2 and adjusted p-value less than 0.05 in at least stimulation.

With this set of 571 cytokine upregulated genes on myeloid cells, we considered if shared cytokine specific responses would dominate over cell-type specific responses to stimulation. Unsupervised clustering and correlation analysis of transcriptional responses showed a clear distinction between stimulations of different cell types. Macrophages and dendritic cells had a more closely correlated response while neutrophils and monocytes were more closely correlated in their response signature **(Fig. 2A)**. Although type 1 (especially IFN-γ and IFN-β) and type 2 (IL-4 and IL-13) cytokine specific responses mainly clustered together within each cell type, this was not sufficient to override the correlation between cell type specific responses. These results indicated that for the most part, the cell type is a larger determinant of whether a gene is upregulated after stimulation than the cytokine. The only exception was a strong correlation between macrophages and dendritic cells stimulated by IFN-β (**Fig. 2A**).

**Figure 2.**
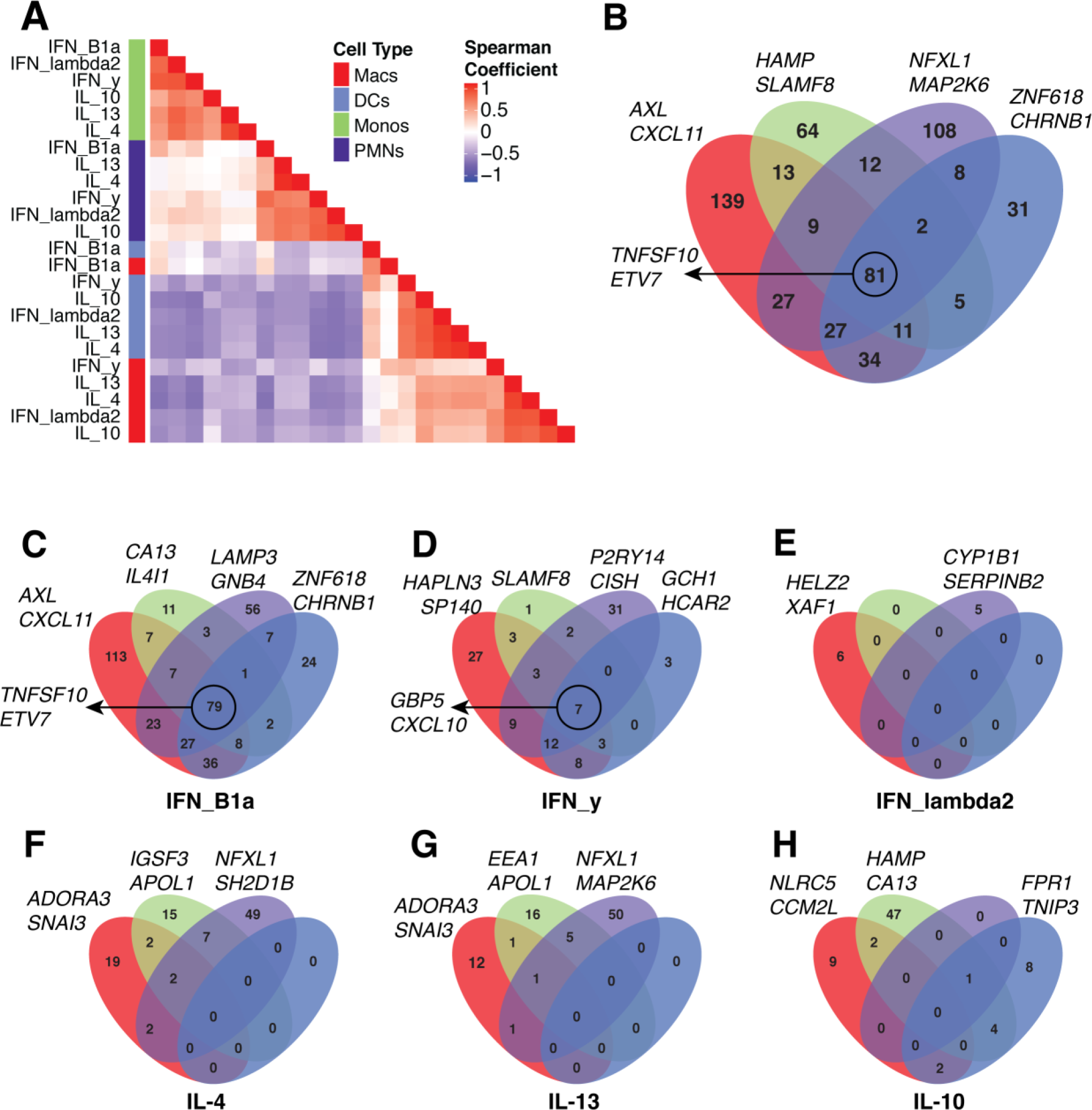
Myeloid cell lineages respond to cytokine stimulation in a cell-type specific manner. (A) Hierarchical clustering of pairwise spearman correlation analysis for the 571 differentially expressed genes. (B) Venn diagrams of 571 genes determined by differential expression in each cell type. 81 of 571 differential genes are shared between all four cell types while 139 (macrophages, red), 64 (monocytes, green), 108 (neutrophils, purple) and 31 (dendritic cells, blue) genes are found to be differentially expressed in only one cell type. (C-H) Venn diagrams for the number of genes significant in each individual cytokine stimulation determined by differential expression in each cell type. The genes listed next to each Venn diagram are the top two differentially expressed genes for each cell type (B), cell type and stimulation (C-H) or the top genes conserved across all four cell types, circled (B-D).

To obtain finer resolution on how the different cell types share responses to cytokine stimulation, we looked for overlaps in differentially expressed genes between cell types. This revealed that 81 of the 571 genes were upregulated in all four cell types **(Fig. 2B)**, which was primarily driven by a shared response to IFN-β stimulation (**Fig. 2C**). However, 342 of the other upregulated genes were specific to a single cell type (**Fig. 2B**), and further segregation by cytokine stimulation confirmed that the major transcriptional response to each cytokine was unique to a particular cell type (**Fig. 2C-H**). For example, IL-10 induced 47 genes that were specific to monocytes, 9 to macrophages and 8 to dendritic cells while having almost no effect on neutrophils (**Fig. 2H**). Alternatively, neutrophils induced 49 and 50 genes uniquely after IL-4 (**Fig. 2F**) and IL-13 (**Fig. 2G**) stimulation while the other cell types were generally less responsive. Neutrophils also had a robust cell type specific response to IFN-γ (31 genes, **Fig. 2D**) and IFN-β stimulation (56 genes, **Fig. 2C**). Overall, these results indicated that the cytokine driven transcriptional responses in different myeloid cell types are highly cell type specific, apart from a core response to IFN-β stimulation (and to a lesser extent IFN-γ) that is shared by all cell types.

### Identification of a myeloid cell cytokine specific transcriptional signature

We next identified specific transcriptional signatures that define a particular cell type and stimulation pair. Through self-organizing map (SOM) analysis (11) we identified clusters of similar gene expression between cytokines in an unbiased manner. For each cell type the full list of differentially expressed genes were sub-clustered into stimulation specific signatures. This analysis divided the gene expression pattern of neutrophils into four sub-clusters corresponding to genes induced only by IFN-β (cluster 1), by both IFN-β and IFN-γ (cluster 2), by both IL-13 and IL-4 (cluster 4) and by IFN-λ (cluster 3) (**Fig. 3A, B**). For macrophages, five clusters were identified corresponding to genes upregulated by only IFN-β (cluster 2), both IFN-β and IFN-γ (cluster 1), both IL-13 and IL-4 (cluster 5), IL-10 (cluster 3) and one cluster which could not be clearly assigned (**Fig. S2A, B**). In dendritic cells, four clusters were identified corresponding to genes upregulated under IFN-β alone (cluster 1), IFN-β and IFN-γ combined (cluster 4), IL10 (cluster 2) and one cluster could not be assigned because two few genes were present (**Fig. S2C, D**). For monocytes, four clusters were identified corresponding to genes upregulated only by IFN-β (cluster 1), both IFN-β and IFN-γ (cluster 3), IL-13 and IL-4 (cluster 4) and IL-10 (cluster 2) (**Fig. S2E, F**). Altogether, 12 cell type and stimulation specific expression patterns could be identified by SOM analysis. Importantly, not all cell types and stimulation signatures were robust enough to be clearly isolated.

**Figure 3.**
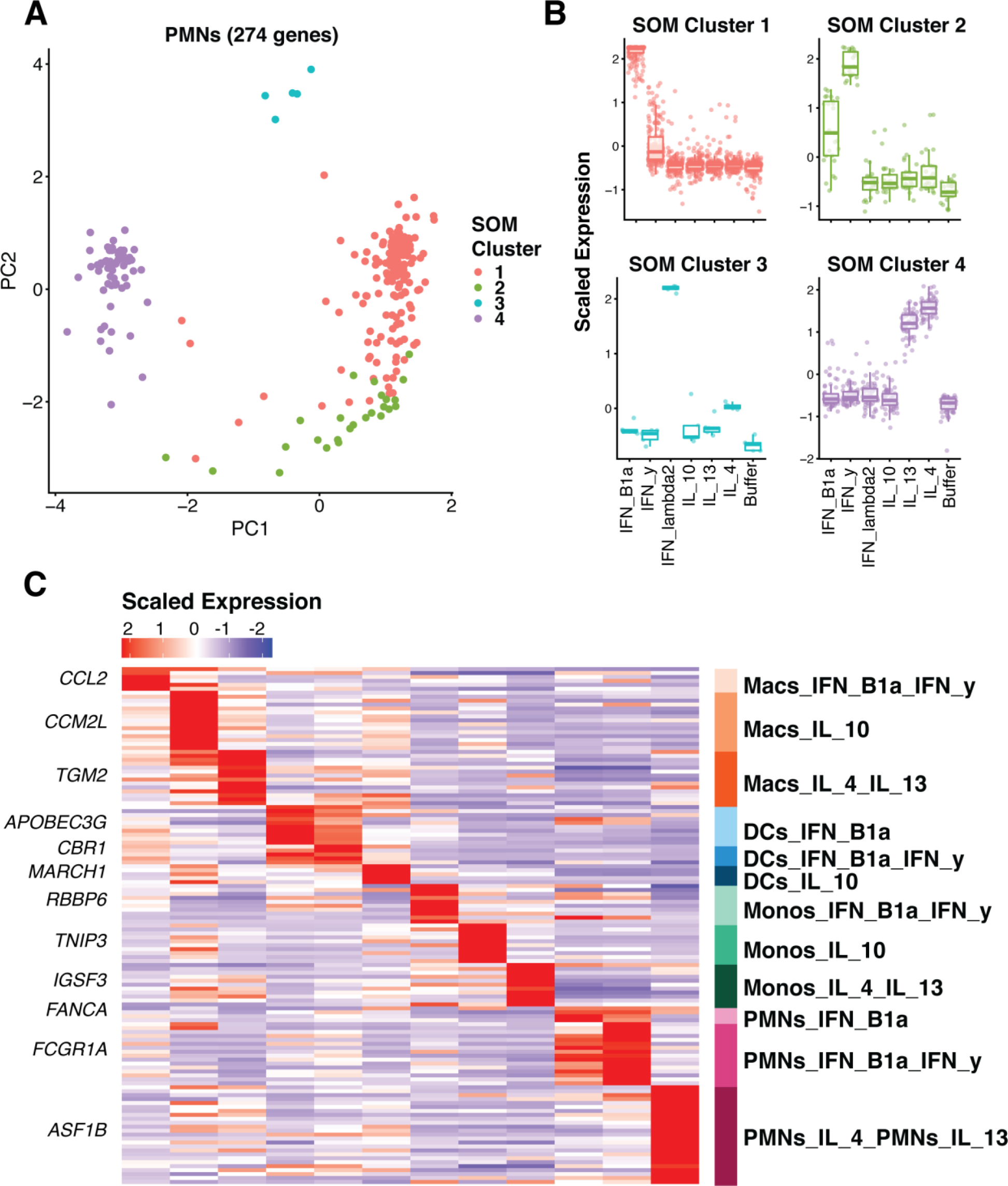
Signature gene expression patterns can be identified in many of the cell types and stimulation conditions. (A) Principle component analysis from self-organizing map (SOM) assignments of gene expression patterns in neutrophils (B). Mapping of SOM clusters by cytokine stimulation. (C) Heatmap indicating the scaled expression levels of selected genes generated from outlier analysis between the three donor samples and between the group assignments derived from SOM analysis. The top gene for each signature is listed.

Following identification of these 12 unique expression clusters, we performed outlier analysis (13) to further filter the expression cluster gene list to only include genes highly specific for the cell type and cytokine stimulation conditions identified by SOM analysis. Genes such as *RBBP6* and *ASF1B* were considered outliers for monocytes responding to IFN-β and IFN-γ and neutrophils responding to IL-4 and IL-13 respectively (**Fig. S3**), due to their highly specific and consistent expression pattern in these cell type stimulation conditions across all three donors. This evaluation identified 131 genes that reflected the 12 myeloid cell cytokine stimulation conditions that were clearly distinguishable (**Fig. 3C, S4 and Table S1**). These genes represent a high confidence marker gene set for myeloid cells under stimulation of various cytokines. We refer to this as a myeloid cell cytokine specific (MCCS) signature.

### Deconvolution of transcriptional signatures from*M. tuberculosis* Infection

To determine the utility of our MCCS signature matrix, we first examined whole-blood transcriptomes from 13 clinical cohorts infected with *M. tuberculosis,* which were publicly available (**Table S2**). Previous studies have described a neutrophil driven type 1 IFN-inducible signature increased in patients with active disease compared to healthy and latently infected individuals (16), hence we were interested in the role of neutrophil specific cytokine responses in this context. More recently, circulating natural killer cells were also reported to increase in abundance during tuberculosis latency but decreased back to baseline during active disease (33). We compiled 8 available human whole blood microarray and 5 RNA-Seq datasets relevant to active tuberculosis infections in GEO and analyzed the two sets independently. We focused our analyses on the differences between healthy (microarray *n* = 88, RNA-Seq *n* = 365), latently infected (microarray *n* = 376, RNA-Seq *n* = 117) and active disease individuals (microarray *n* = 547, RNA-Seq *n* = 306) as described in **Table S2**. We first utilized the original LM22 basis matrix from CIBERSORT (https://cibersort.stanford.edu)(1) and the more recent ‘immunoStates’ matrix (15) to infer leukocyte representation by support vector regression through CIBERSORT. The original LM22 basis matrix identifies 22 human hematopoietic cell phenotypes from peripheral blood and *in vitro* culture conditions while immunoStates identifies 20 immune cell types from over 6,000 samples during different disease states. Using these matrices, we were able to confirm that CD56bright NK cells (immunoStates) were increased in abundance for latently infected individuals both in the microarray and RNA-Seq datasets (**Fig. 4A**). While the signature of resting NK cells (LM22) also showed this response (**Fig. S5E**) in the microarray dataset, the RNA-Seq dataset showed a slightly different pattern (**Fig. S5F**). This finding is consistent with immunoStates being an improved basis matrix compared to LM22 and confirmed that our compiled datasets could reproduce previously published findings (33).

**Figure 4.**
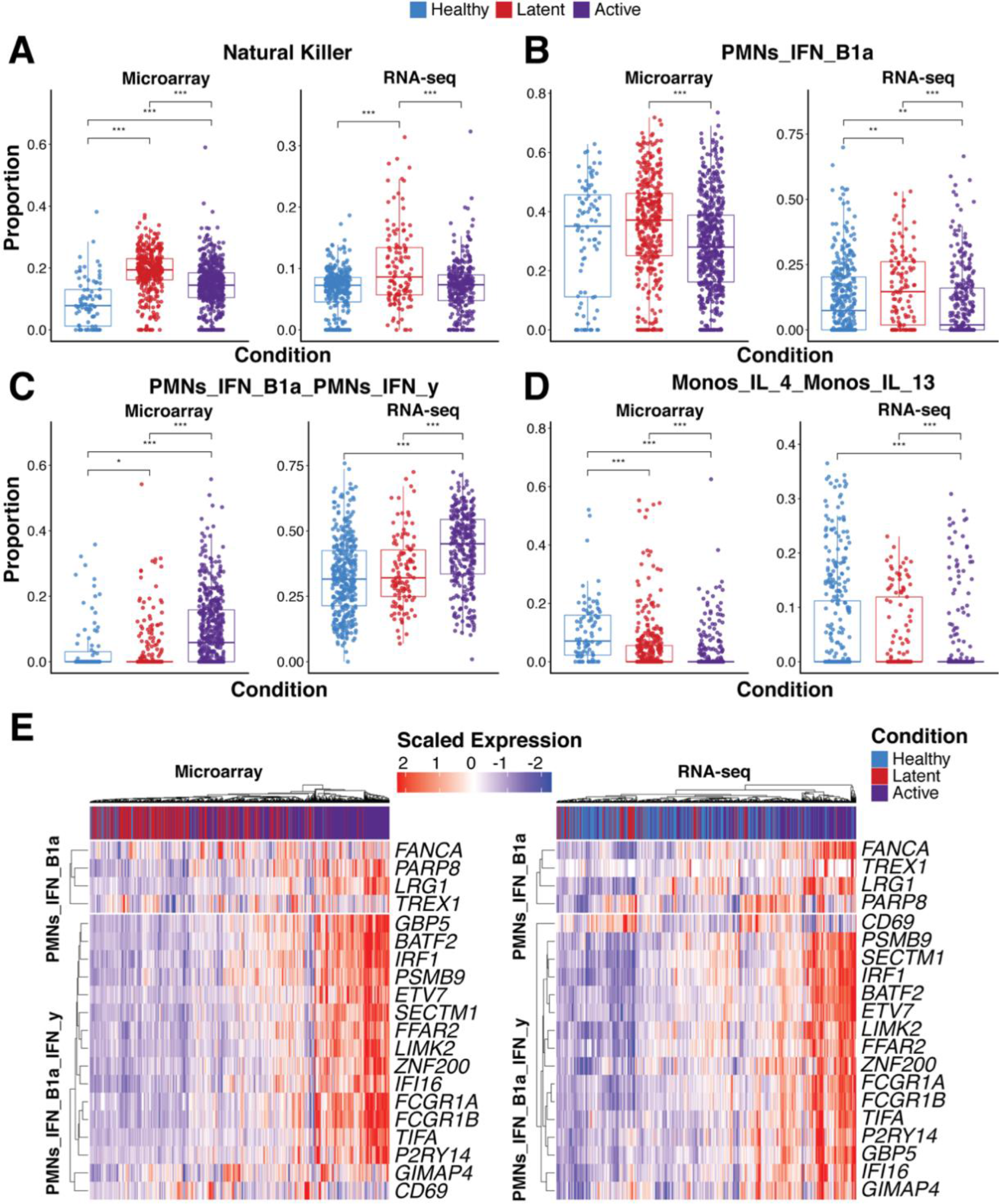
Statistical deconvolution of bulk expression profiles indicates role of interferon-induced neutrophil response *in M. tuberculosis* infection. (A-D) Proportion estimates for neutrophils, Monocytes and natural killer (NK) cells from CIBERSORT with our MCCS signature matrix (B-D) and immunoStates (A) for 8 microarray datasets and 5 RNA-Seq datasets (Table S2). (E) Scaled expression of 20 genes found in our neutrophil-interferon signatures are shown for the RNA-Seq and microarray samples as well as the disease status of the sample. Significance was determined by Kruskal-Wallis rank sum test with *p*-value < 0.05 = *, *p*-value < 0.01 = ** and *p*-value < 0.001 = ***. Sample sizes for each disease state and data type are as follows; healthy (microarray *n* = 88, RNA-Seq *n* = 365), latently infected (microarray *n* = 376, RNA-Seq *n* = 117) and active disease individuals (microarray *n* = 547, RNA-Seq *n* = 306).

When we examined the inferred abundance of neutrophils, we found that the LM22 matrix indicated an increased abundance of neutrophils in actively infected individuals from the microarray dataset (**Fig. S5E)**, but also suggested that neutrophils were more abundant in latently infected individuals compared to healthy individuals from the RNA-Seq dataset (**Fig. S5F)**. In contrast, the immunoStates matrix inferred greater abundance of neutrophils during active disease from the RNA-Seq dataset (**Fig. S5D)** with decreased abundance of neutrophils during latent infection in the microarray dataset (**Fig. S5C)**. When we applied our MCCS matrix on these datasets, we found that there was a clear increase in actively infected individuals for neutrophil response genes that were inducible by both IFN- γ and IFN- β (**Fig. 4C**). Surprisingly, genes that were only inducible by IFN- β in neutrophils were reduced in expression during active infection compared to latent infection (**Fig. 4B**). This was consistent for both microarray and RNA-Seq datasets. Although a role for IFN- β during active tuberculosis infection has now been well established (16), these results were surprising in that they point to a requirement for both IFN-γ and IFN- β in driving the IFN-inducible signature of neutrophils during active tuberculosis. Alternatively, it is perhaps impossible to truly determine if the IFN-inducible signature of neutrophils is the result of type 1 or type 2 IFNs since they induce a similar set of genes (34). Notably, when we examined other myeloid cell responses, we found that there was a consistent reduction of the IL-4/IL-13 signatures from both monocytes (**Fig. 4D**) and macrophages (**Fig. S5A,B**) during active infection, relative to healthy and latently infected individuals. Hence, in addition to providing further insights into the IFN-inducible neutrophil signature during human tuberculosis, our MCCS matrix implicates a suppression of type-2 cytokine (IL-4 and IL-13) responses in monocytes and macrophages during active infection. Additionally, there was an increased abundance of dendritic cells (DCs) expressing IFN- γ and IFN-β inducible genes during active infection (**Fig. S5A,B**). From these results, we were able to gain additional biological insight into the cytokine responses of myeloid cells during different stages of tuberculosis infection.

### Interleukin-stimulated Neutrophil Signature Indicates Poor Survival in Glioma

Recently, infiltrating and circulating myeloid cells have been tied to survival and likelihood of response to immunotherapy in the context of human gliomas (35, 36). A significant portion of the cellular mass in primary glioma samples is infiltrating immune cells such as tumor-associated macrophages (TAMs), whose levels correlate with tumor grade and severity, and other myeloid subsets (37). Additionally, over 600 primary glioma tumors have been profiled by the Cancer Genome Atlas (TCGA) (31) by a variety of sequencing methods including RNA-Seq with detailed clinical outcome information. Applying statistical deconvolution based on our curated MCCS signature, we found a strong but reciprocal relationship to survival for neutrophils responding to IL-4 and IL-13 stimulation, and monocytes responding to IFN-β and IFN-γ stimulation. Monocyte IFN responses were predictive of favorable survival, whereas tumors with high neutrophil IL-4/IL-13 responses exhibited reduced patient survival (**Fig. 5A and S6**).

**Figure 5.**
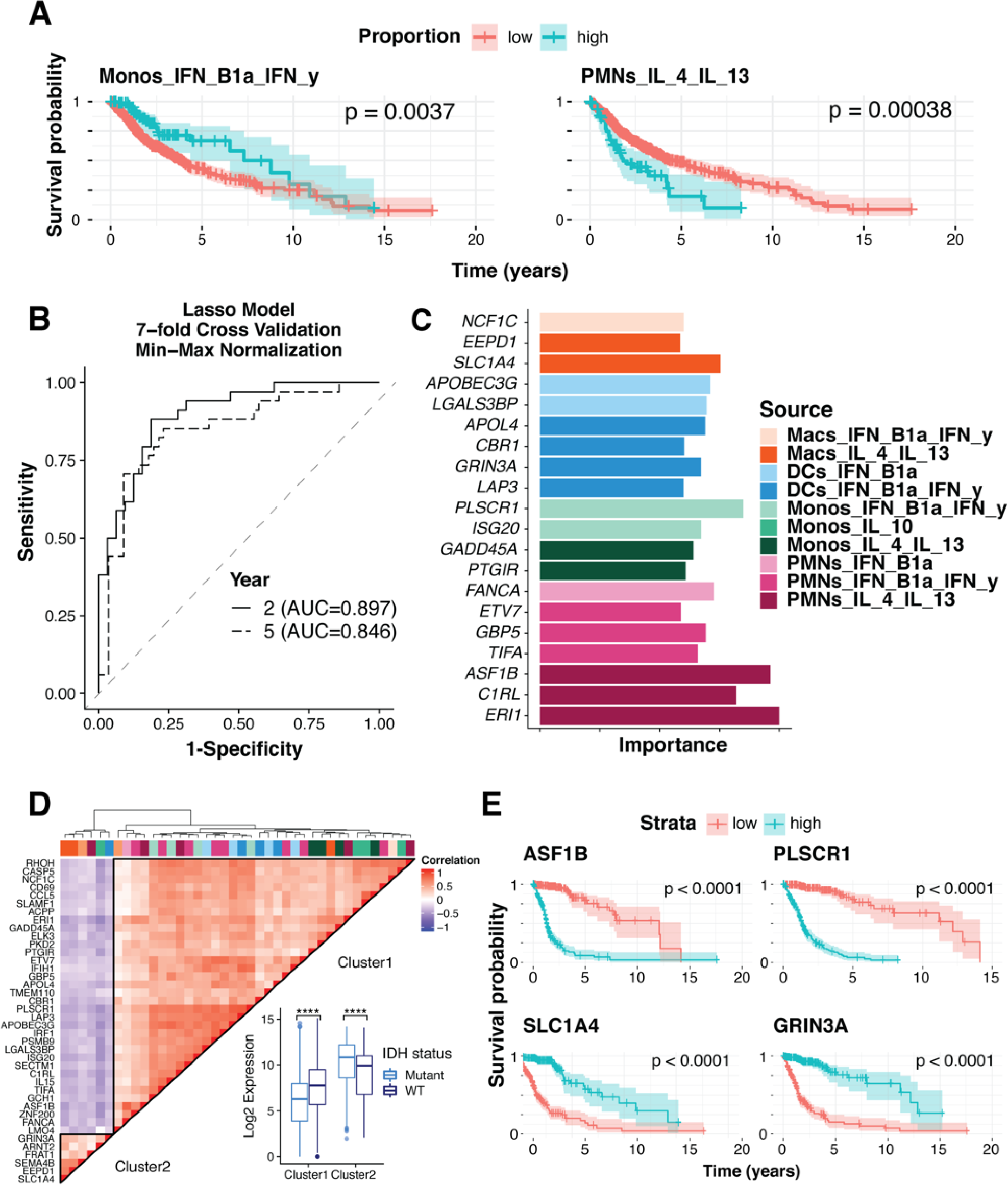
Myeloid signatures under stimulation are indicative of survival in glioma. (A) Survival analysis of statistically deconvolved bulk RNA-Seq data from 671 glioma tumor samples for individuals with low proportion estimates (red) or high proportion estimates (blue) for Neutrophils responding to IL4 and IL13 (PMNs_IL4_IL13) and for Monocytes responding to IFN- γ and IFN- β (Monos_IFN_B1a_IFN_y). (B) The power of our myeloid gene signature was determined by area under the curve measures for LASSO models at 2 and 5-year increments trained on our 131-cytokine stimulated myeloid gene signature with 7-fold cross validation. A dashed diagonal line indicates an AUC of 0.5 for a random prediction model. (C) As measured by model importance (see methods) the top 20 features derived from the 5-year prediction model are shown. (D) Hierarchical clustering of pairwise spearman correlation analysis of 40 of the most predictive features derived from our 5-year model. Gene expression clusters were then mapped by genotype for a wild-type or mutated IDH gene locus, a molecular marker of gliomas. (E) Survival analysis based on individual genes from cluster 1 (ASF1B and PLSCR1) and cluster 2 (SLC1A4 and GRINA3A) using a cox regression model of gene expression in the TCGA samples profiled.

We next considered a more direct approach to assess the utility of our MCCS signature to predict survival of patients with glioma. We trained least absolute shrinkage and selection operator (LASSO) models on our 131-gene MCCS signature, the original LM22 (1) basis matrix and the immunoStates (15) basis matrix separately to classify 2 and 5-year survival predictions. Our model demonstrated robust survival prediction with an area under the ROC curve (AUC) between 0.85 (5-year) and 0.89 (2-year) on our test set while the LM22 and immunoStates signatures were lower (immunoStates AUC = 0.868 at 2 years and 0.763 at 5 years, LM22 AUC = 0.828 at 2 years and 0.788 at 5 years) (**Fig 5B and S7**). Evaluation of the gene importance for survival predictions in our MCCS matrix at 5 years indicates that the top genes were derived from the IL-4/IL-13-stimulated neutrophils and IFN-β and IFN-γ-stimulated monocytes (**Fig. 5C**), confirming the CIBERSORT proportion estimates and survival curves shown in **Figure 5A**. In addition to the cell type and stimulation condition we were also interested in the relationships between the genes most predictive of long-term survival. Correlation analysis of the top features with strong predictive power, as measured by feature importance (See methods) indicated two distinct expression profiles (**Fig 5D**). Furthermore, primary glioma samples from TCGA have been previously profiled to identify somatic mutations and molecular markers (31) indicative of survival. One such marker is the gene encoding isocitrate dehydrogenase (IDH), which when mutated is known to be associated with increased patient survival in both low and high-grade gliomas (38). Based on pairwise gene expression correlation analysis of the 40 most predictive gene features from our model, we identified two clusters which were found to significantly differ in their gene expression between glioma samples with a mutated or wild type IDH gene (**Fig 5D**). Specifically, on average cluster 1 genes had higher expression in samples with wild type IDH status while cluster 2 genes have significantly higher expression in samples with a mutated IDH gene. This indicated that our set of genes were not only predictive of survival but also strongly associated with known molecular markers for primary gliomas.

Given the strength of the importance measures for several of the top features we also measured survival outcomes based on gene expression levels with a cox regression for *ASF1B, PLSCR1, SLC1A4* and *GRIN3A* and found significant associations between these expression-based models and survival (**Fig. 5E**). *ASF1B* and *PLSCR1* gene expression were indicative of poorer survival outcomes while *SLC1A4* and *GRIN3A* expression were indicative of more favorable outcomes (**Fig 5D, E**). Further, *ASF1B*, a strong indicator of glioma prognosis, was derived from the neutrophil signature in response to IL-13 and IL-4 suggesting a more complex role for neutrophils in the tumor microenvironment. Interestingly, expression of *SLC1A4*, identified as part of the IL-4 and IL-13 stimulated macrophage signature, was indicative of better survival (**Fig 5E**) raising additional questions about the role of TAMs in primary glioma samples. Altogether, our MCCS signature matrix was able successfully predict patient survival from gene expression in primary glioma samples corresponding to specific neutrophil-associated gene signatures and other myeloid cell signatures.

## DISCUSSION

In this study, we first assessed the transcriptional response of 4 different human myeloid cell types to stimulation with a panel of cytokines. This enabled us to assemble a set of gene signatures for myeloid cell type–cytokine specific response genes, which we could then assess for biological and clinical relevance. Although limited to neutrophils, monocytes, macrophages and dendritic cells currently, the signature matrix provides the cellular context these cells experience during cytokine stimulation. This approach could be expanded to include additional cell types as well as additional stimulation conditions to provide even more granular context. Hence, controlled *in-vitro* assays could be quite relevant towards interpreting the expression profiles *in vivo* for primary human blood and tissue samples. This approach can thus be applied towards existing bulk transcriptomics data available in GEO, for example from GTeX and TCGA.

In the context of *M. tuberculosis* infection the importance of an interferon-inducible gene signature is well documented (39). The first seminal study, which also profiled purified cell populations had indicated that this signature was driven by neutrophils and both IFN-γ and type I interferon signaling (16). Our findings here are consistent with that initial report, since actively infected individuals were enriched for neutrophil response genes that are inducible by both IFN-γ and IFN-β (**Fig. 4C**). However, we found that neutrophil genes inducible by IFN-β alone are reduced in actively infected individuals indicating that IFN-γ may be more dominant than type 1 interferons in driving the interferon-inducible signature of neutrophils during active tuberculosis. This is in contrast to a recent report showing that *IFNG* (which encodes IFN-γ) and *TBX21* (which encodes the transcription factor T-bet) are downregulated in patients with active TB (17). Hence, the ratio of type 1 interferon vs IFN-γ inducible genes in neutrophils needs to be better clarified in future studies. Since the goal of our study was to explore the biological context of myeloid cells responding to cytokine stimulation, rather than to identify the ideal gene signature for discriminating active TB from latent TB, we have not performed deeper characterization of heterogeneity in the multiple datasets that we compiled from TB patients.

The relationship between neutrophil responses to IL-4 and IL-13 stimulation with glioma survival was of particular interest. Previous reports from helminth infected mice have described a distinct transcriptional response to type 2 cytokines in neutrophils (40) and the concept of N2 neutrophils in the tumor microenvironment has also been proposed (41, 42). However, the transcriptional responses of human neutrophils to stimulation by IL-4 and IL-13 has not been well established. Instead, TGF-β has been implicated in N2 polarization (43), which was not examined as part of our analysis. Our results demonstrate not only that human neutrophils respond to IL-4 and IL-13 stimulation with a very distinct transcriptional signature but also that this signature can be detected in tumor samples and is associated with survival outcomes for glioma in particular. Therefore, we provide some of the best evidence thus far that type-2 cytokine associated neutrophil activation may play an important role in tumor progression.

An important limitation of our study is that transcripts that were found to be associated with specific myeloid cell type-cytokine stimulation combinations could also be expressed by other immune or non-immune cells. While we are inferring or interpreting some of these results in the context of myeloid cell responses, the same transcripts could be induced by other cell types in response to other cytokines we have not examined. Future studies should expand upon this preliminary assessment of 4 myeloid cell types and 6 cytokine combinations, to include multiple immune and non-immune cell types and additional cytokines or other micro environmental stimuli. Additionally, we have not assessed combinations of cytokines at varying concentrations. In an inflamed environment, a combination of different cytokines at different concentrations will have synergistic or inhibitory effects on different cell populations.

Recently, approaches have been developed to utilize single-cell transcriptomics data for deconvolution of bulk transcriptomic data. While this approach could in principle assess hundreds or thousands of cell states in bulk transcriptomic data, the reference collection sample set for the scRNA-Seq profiles may not provide easily interpretable data on the cytokine environment of the bulk tissue. We are currently working towards combining specific cytokine stimulation conditions and scRNA-Seq to determine if we can assemble a cytokine specific matrix for hundreds or thousands of single cell states.

We present here the concept of combining transcriptional profiles from *in vitro* stimulated immune cells with different cytokines, together with algorithms such as CIBERSORT (1) to infer the cytokine and immune cell environment within an inflamed tissue. We also provide a myeloid cell cytokine signature matrix that can be used by the community to help assess immune cell composition in complex samples. This approach has the potential to provide additional biological insights into the ever-expanding collections of transcriptional profiling datasets associated with different diseases, potentially leading to improvements in diagnosis and therapeutic strategies during infection and tumor progression.

## ACKNOWLEDGEMENTS

We would like to thank Evelien T. M. Berends for help setting up the primary human myeloid cell system in the Torres lab. We would like to acknowledge The Genome Technology Center, a shared resource partially supported by the Cancer Center Support Grant P30CA016087 at the Laura and Isaac Perlmutter Cancer Center, for preparing and sequencing the RNA libraries. We would also like to thank Dr. Matija Snuderl for his guidance and expertise in gliomas.

## Footnotes

1 **FUNDING** Research in the V.J.T. lab and P.L lab is supported in part by the National Institutes of Health under award numbers AI099394, AI121244, AI105129, AI130945, AI133977, DK103788, HL084312 and the Department of Defense (W81XWH-16-1-0256). E.E.Z. was supported by an NIAID-supported institutional research training grant on Infectious Disease & Basic Microbiological Mechanisms T32AI007180. The NYU Langone Health Genome Technology Center is a shared resource partially supported by the Laura and Isaac Perlmutter Cancer Center, Cancer Center Support Grant P30CA016087. V.J.T. is a Burroughs Wellcome Fund Investigator in the Pathogenesis of Infectious Diseases. The funders had no role in study design, data collection and interpretation, or the decision to submit the work for publication.

