## Supplemental Figures for "Distinct features of human myeloid cell cytokine response profiles identify neutrophil activation by cytokines as a prognostic feature during tuberculosis and cancer1"

### SUPPLEMENTARY TABLES

**Supplementary Table 1. Myeloid cell cytokine specific signature matrix.**

MCCS table of the averaged gene expression values for myeloid cell cytokine specific signatures across all three donors. Colors indicate the specific cell type and cytokine stimulation condition.

**Supplementary Table 2. Description of samples and cohorts of *M. tuberculosis* infection.**

Sample breakdown and number of samples from RNA-Seq and Microarray as well as sample stratification (healthy, latent or active infection status) are included. A full list of the sample IDs obtained from GEO for GSE19491, GSE28623, GSE37250, GSE39939, GSE39940, GSE40553, GSE41055, GSE56153, GSE101705, GSE107995, GSE79362, GSE89403 and GSE94438 are included.

**SUPPLEMENTARY FIGURES**

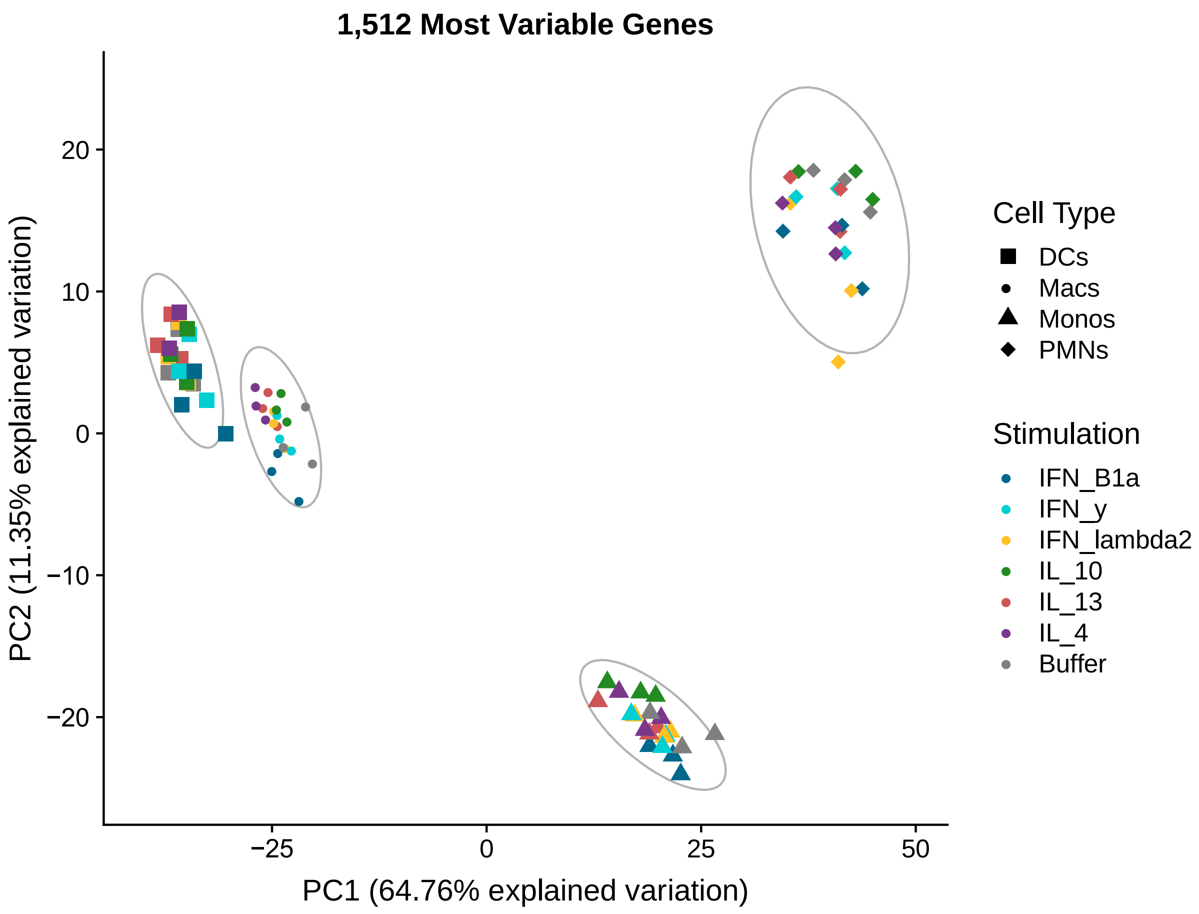


**Figure S1. Cell type specific differences are greater than donor differences.** Principle component analysis of 1,512 variably expressed genes determined by a 90% quantile filter for variance across all cell types and stimulations including buffer. Cell types are indicated by the corresponding symbol while stimulation is colored as indicated.

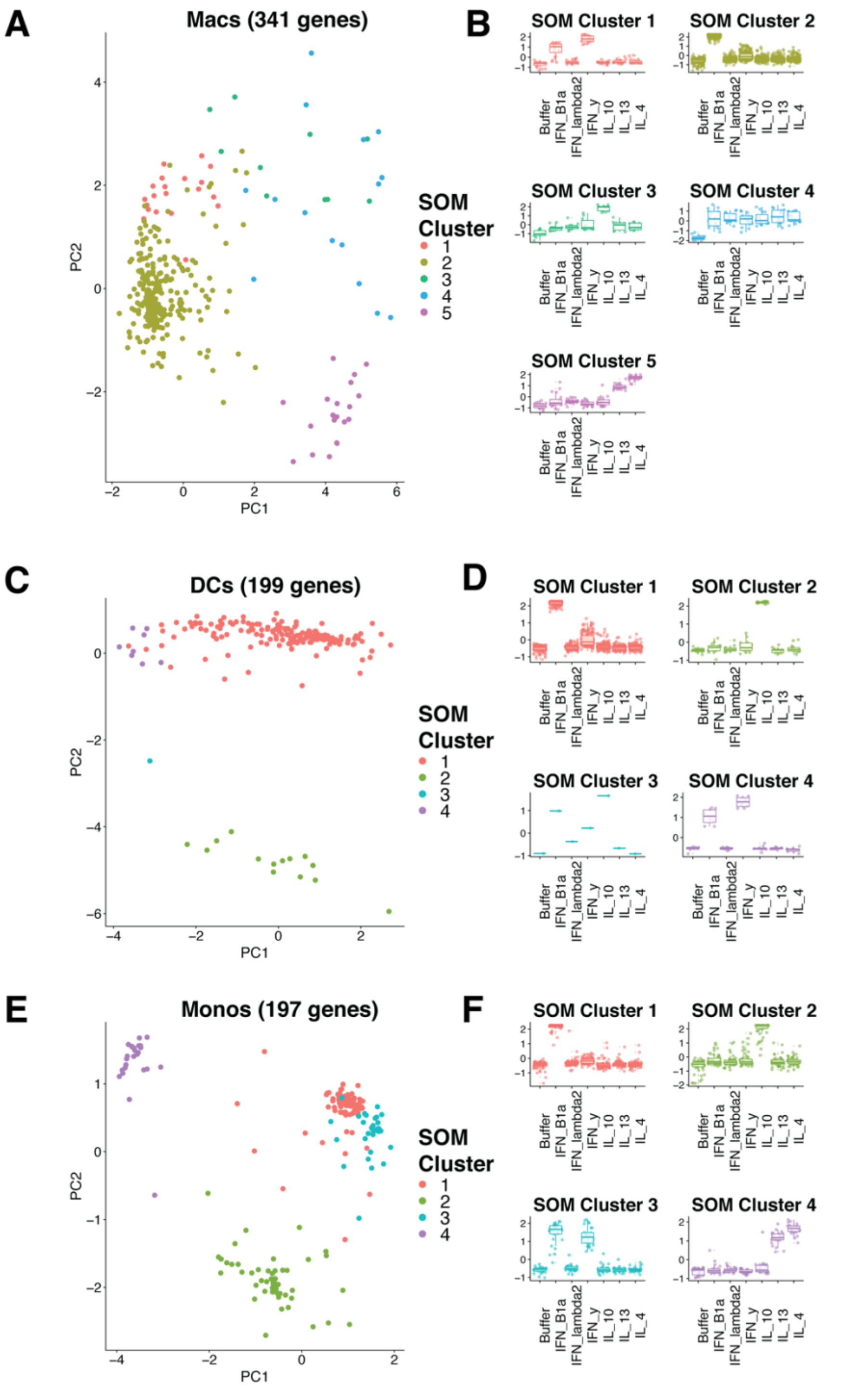


**Figure S2. Specific cytokine stimulations share expression patterns within cell types.** Self-organizing map analysis for cell type and stimulation clustering in Macrophages (A, B), dendritic cells (C, D), and Monocytes (E, F). Principle component analysis of each differentially expressed gene is shown to determine underlying clusters (A, C, E). Scaled gene expression levels for stimulations in each cluster for each cell type (B, D, F).

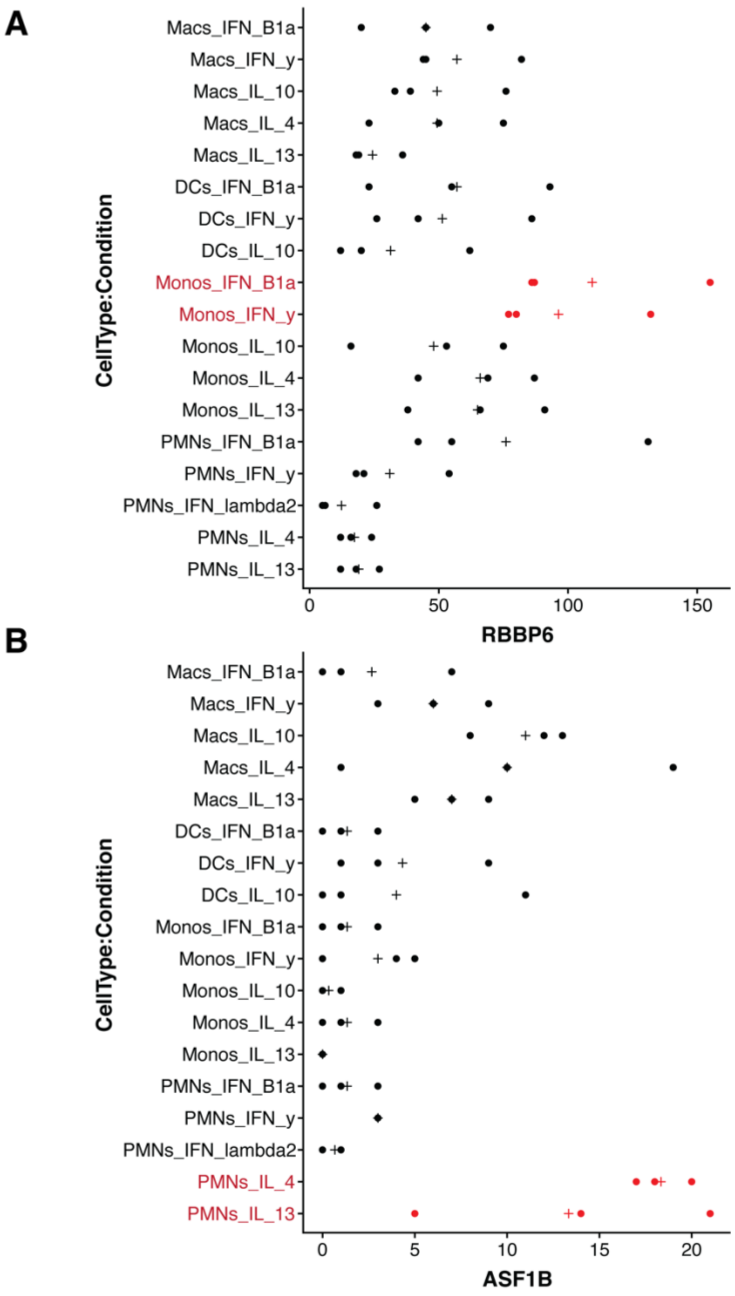


**Figure S3. Outlier analysis confirms high-confidence genes within each cell type and stimulation condition.** RBBP6 (A) and ASF1B (B) are shown as representative gene signatures in IFN-β and IFN-γ stimulated monocytes and IL4 and IL13 stimulated neutrophils respectively. Each dot represents one of three donors and the '+' symbol indicates the mean expression level.

A

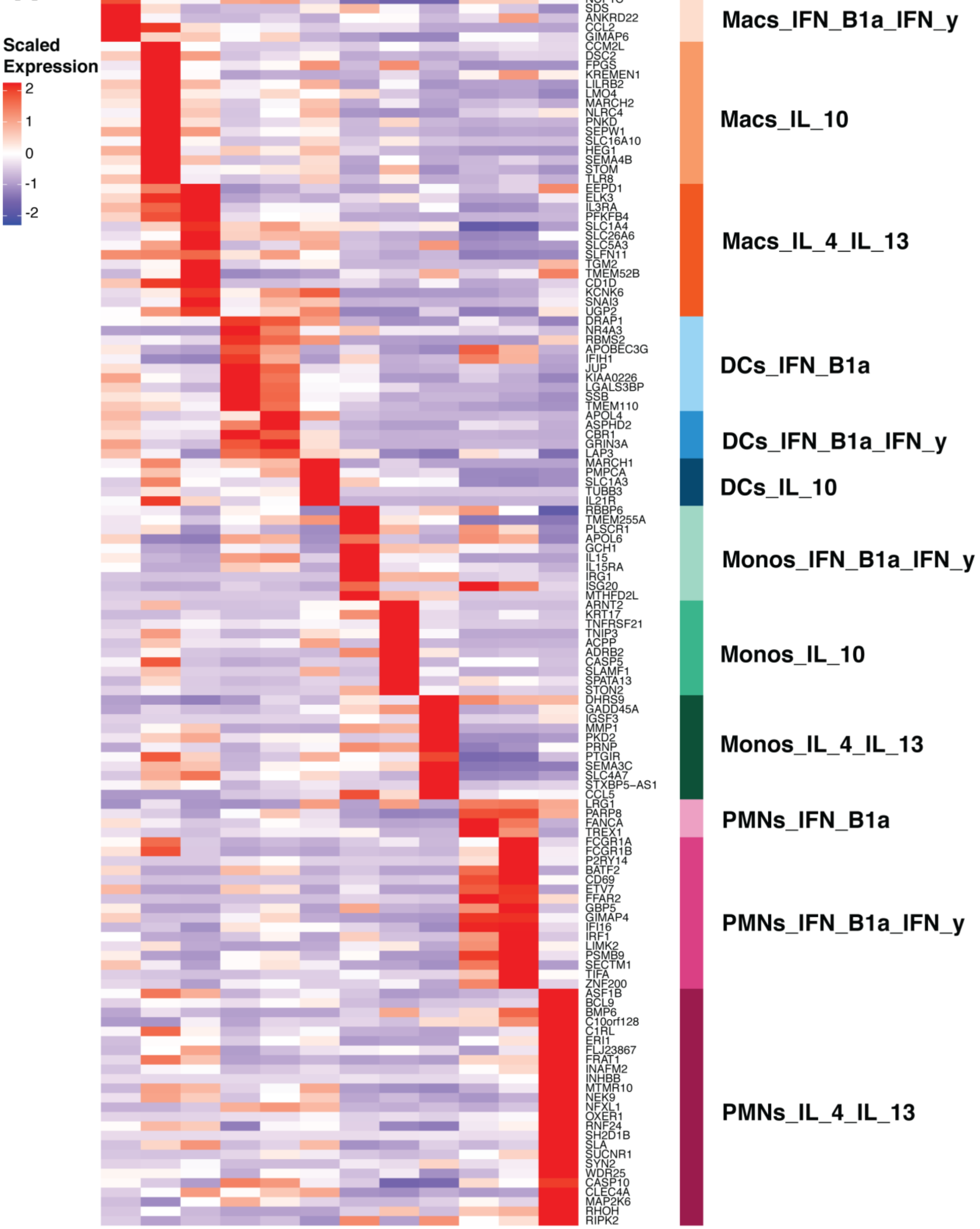

Scaled Expression
2
1
0
-1
-2
GCNT2
NCF1C
SDS
ANKRD22
CCL2
GIMAP6
CCM2L
DSC2
FPGS
KREMEN1
LILRB2
LMO4
MARCH2
NLRC4
PNKD
SEPW1
SLC16A10
HEG1
SEMA4B
STOM
TLR8
EEPD1
ELK3
IL3RA
PFKFB4
SLC1A4
SLC26A6
SLC5A3
SLFN11
TGM2
TMEM52B
CD1D
KCNK6
SNAI3
UGP2
DRAP1
NR4A3
RBMS2
APOBEC3G
IFIH1
JUP
KIAA0226
LGALS3BP
SSB
TMEM110
APOL4
ASPHD2
CBR1
GRIN3A
LAP3
MARCH1
PMPCA
SLC1A3
TUBB3
IL21R
RBBP6
TMEM255A
PLSCR1
APOL6
GCH1
IL15
IL15RA
IRG1
ISG20
MTHFD2L
ARNT2
KRT17
TNFRSF21
TNIP3
ACPP
ADRB2
CASP5
SLAMF1
SPATA13
STON2
DHRS9
GADD45A
IGSF3
MMP1
PKD2
PRNP
PTGIR
SEMA3C
SLC4A7
STXBP5-AS1
CCL5
LRG1
PARP8
FANCA
TREX1
FCGR1A
FCGR1B
P2RY14
BATF2
CD69
ETV7
FFAR2
GBP5
GIMAP4
IFI16
IRF1
LIMK2
PSMB9
SECTM1
TIFA
ZNF200
ASF1B
BCL9
BMP6
C10orf128
C1RL
ERI1
FLJ23867
FRAT1
INAFM2
INHBB
MTMR10
NEK9
NFXL1
OXER1
RNF24
SH2D1B
SLA
SUCNR1
SYN2
WDR25
CASP10
CLEC4A
MAP2K6
RHOH
RIPK2
Macs_IFN_B1a_IFN_y
Macs_IL_10
Macs_IL_4_IL_13
DCs_IFN_B1a
DCs_IFN_B1a_IFN_y
DCs_IL_10
Monos_IFN_B1a_IFN_y
Monos_IL_10
Monos_IL_4_IL_13
PMNs_IFN_B1a
PMNs_IFN_B1a_IFN_y
PMNs_IL_4_IL_13

**Figure S4 Myeloid cell cytokine stimulation signature matrix.** 131 myeloid cell cytokine stimulation (MCCS) signature matrix with gene names.

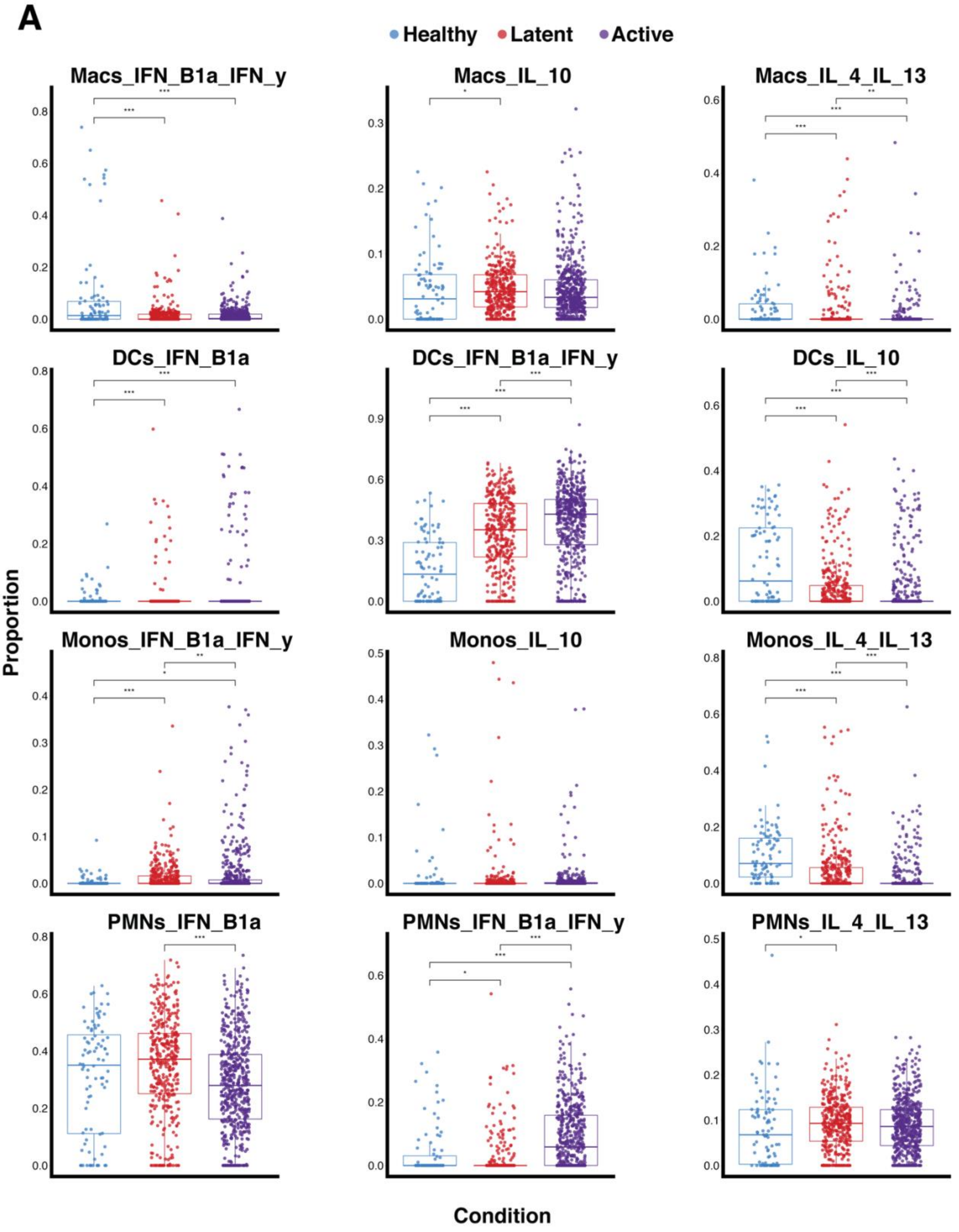


Matrix: MCCS
Data Type: Microarray

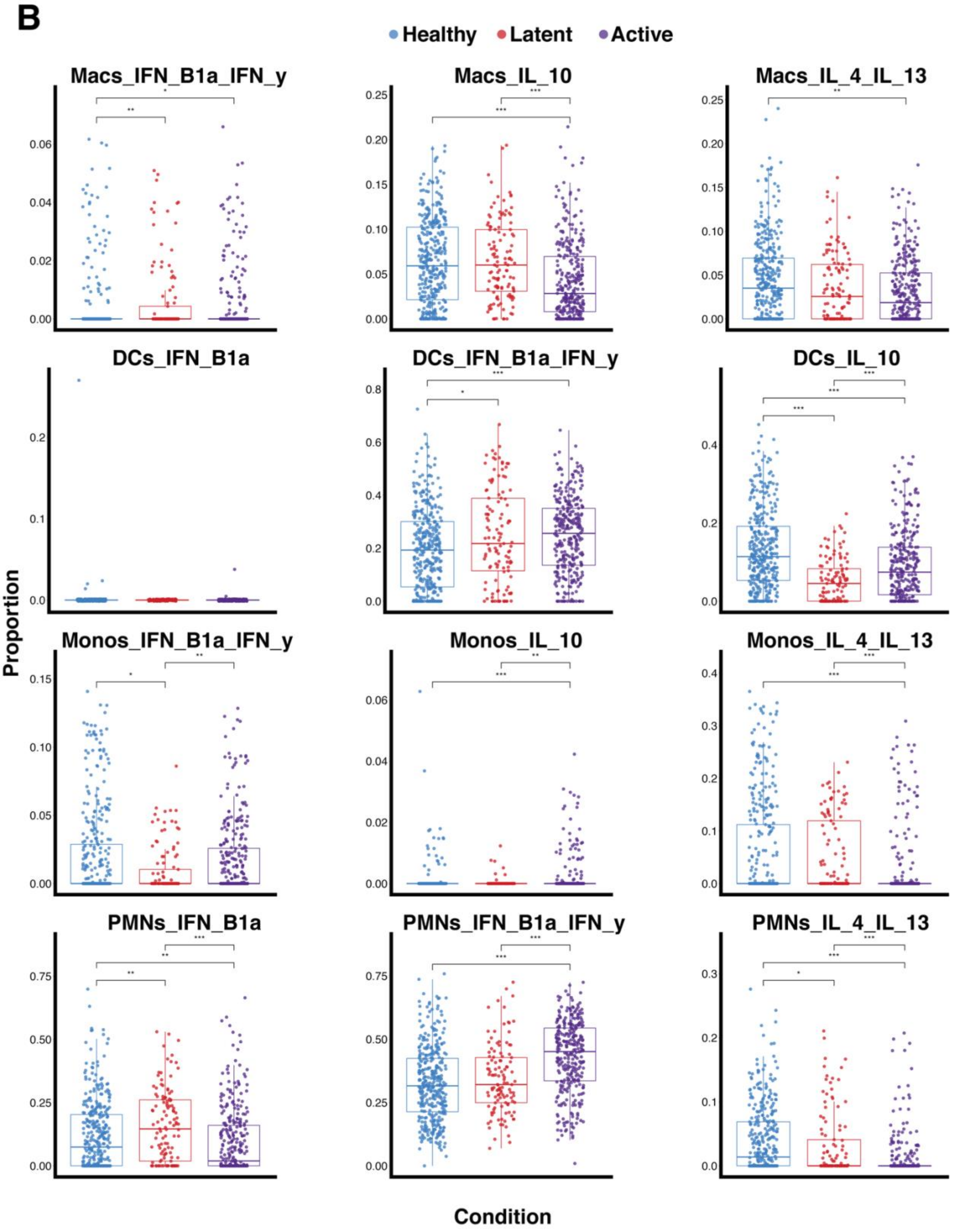


**Matrix: MCCS**
**Data Type: RNA-Seq**

C

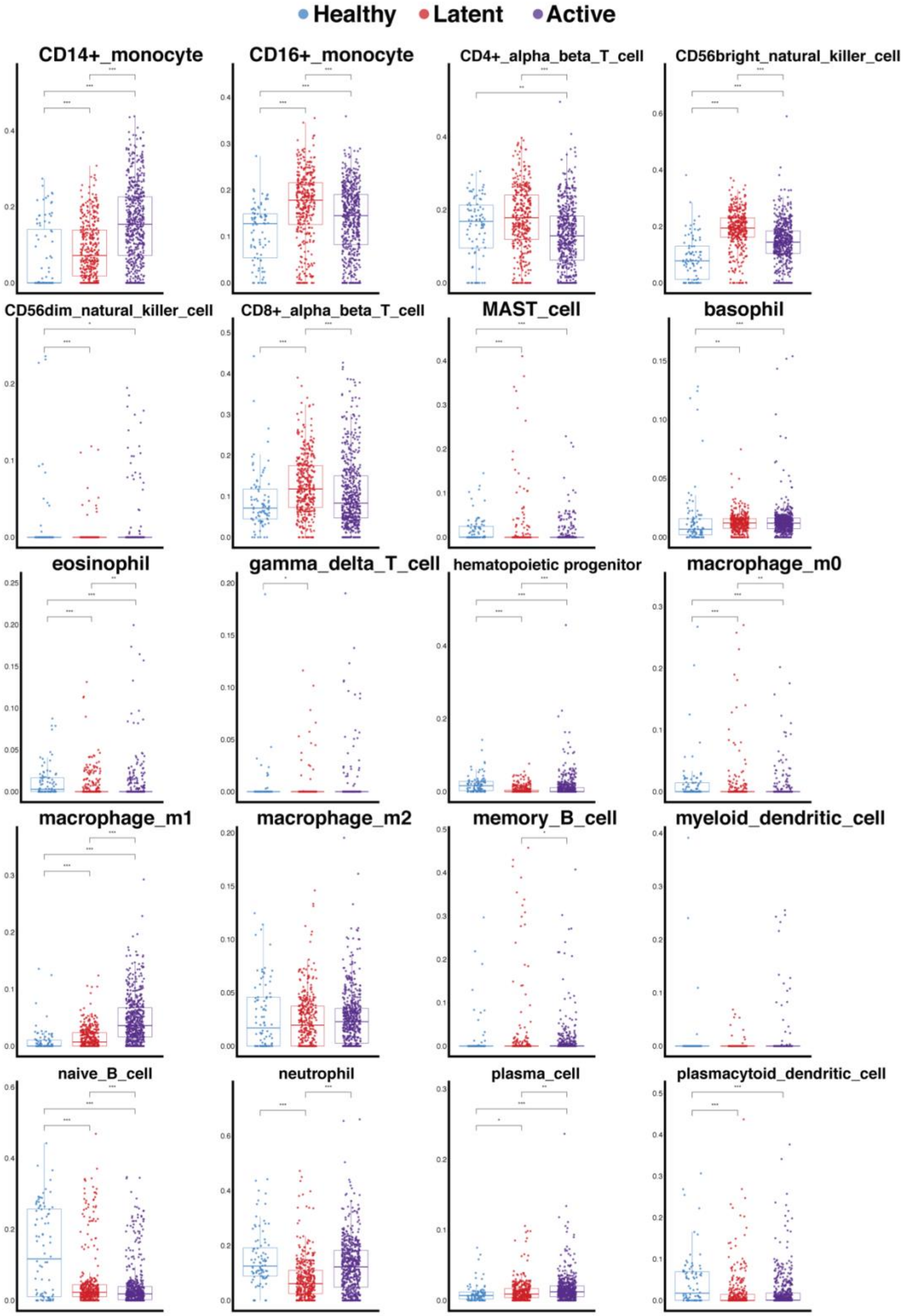


**Matrix: ImmunoStates**
**Data Type: Microarray**

D

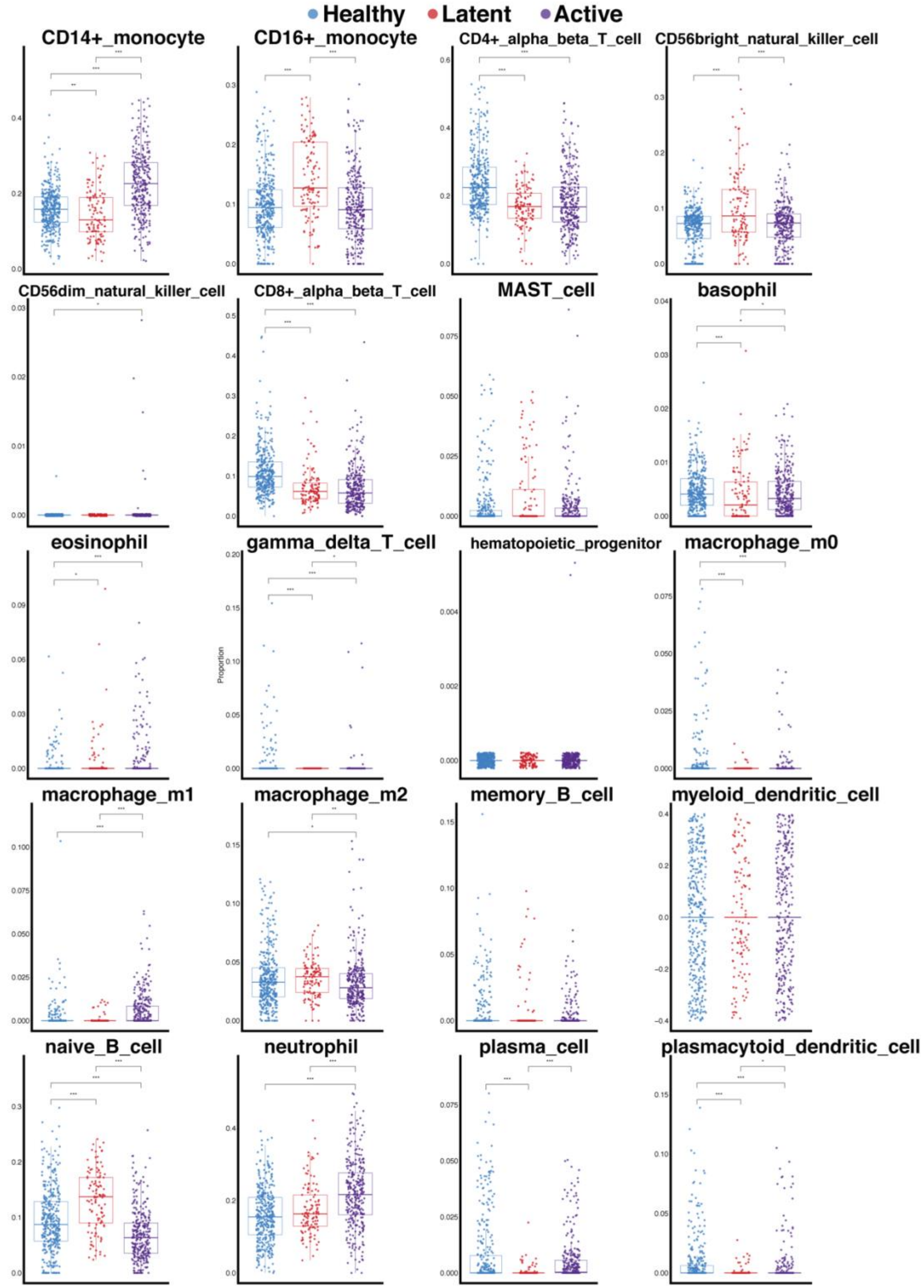


**Matrix: ImmunoStates**
**Data Type: RNA-Seq**

**E**

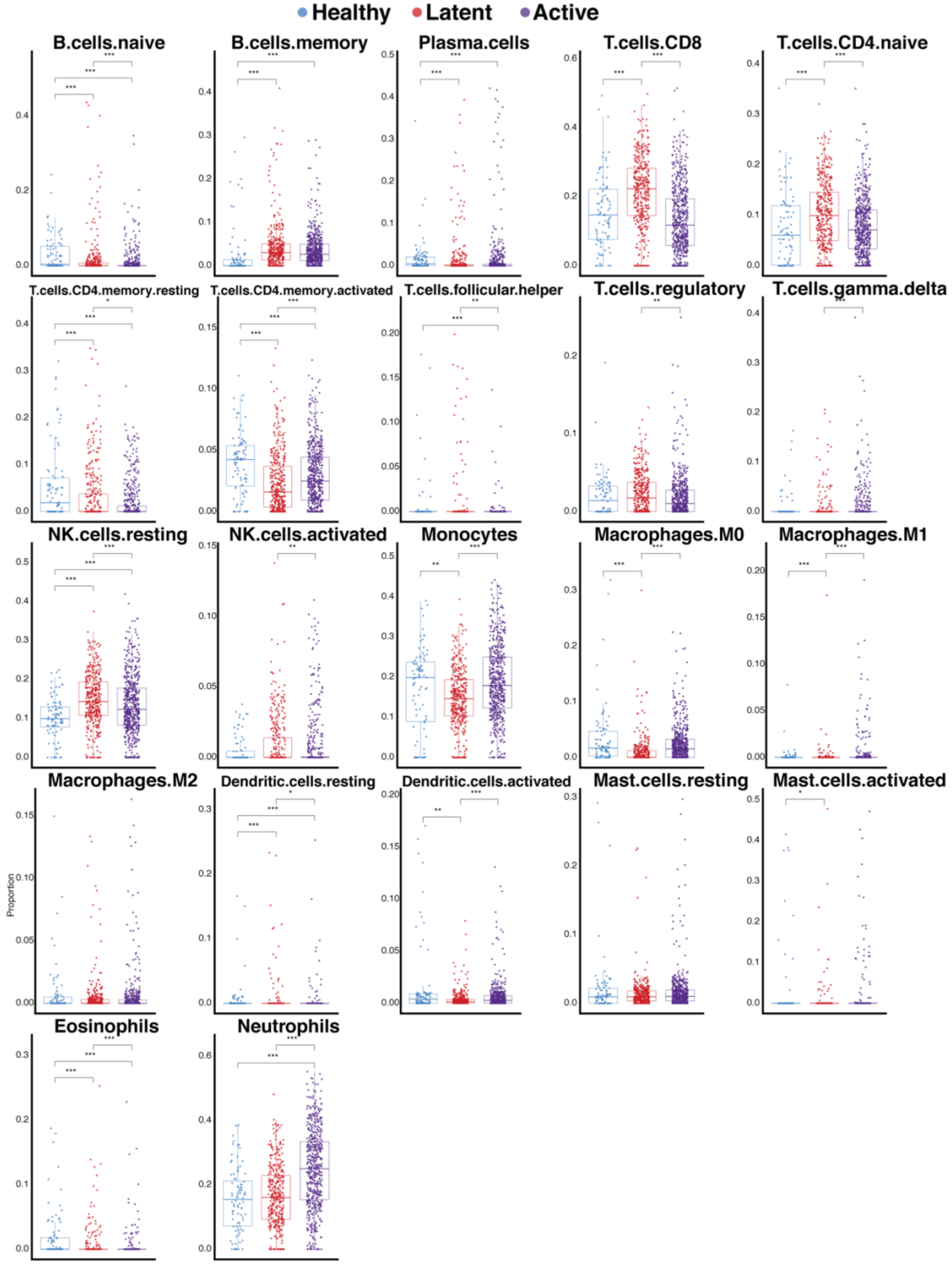


**Matrix: LM22**
**Data Type: Microarray**

F

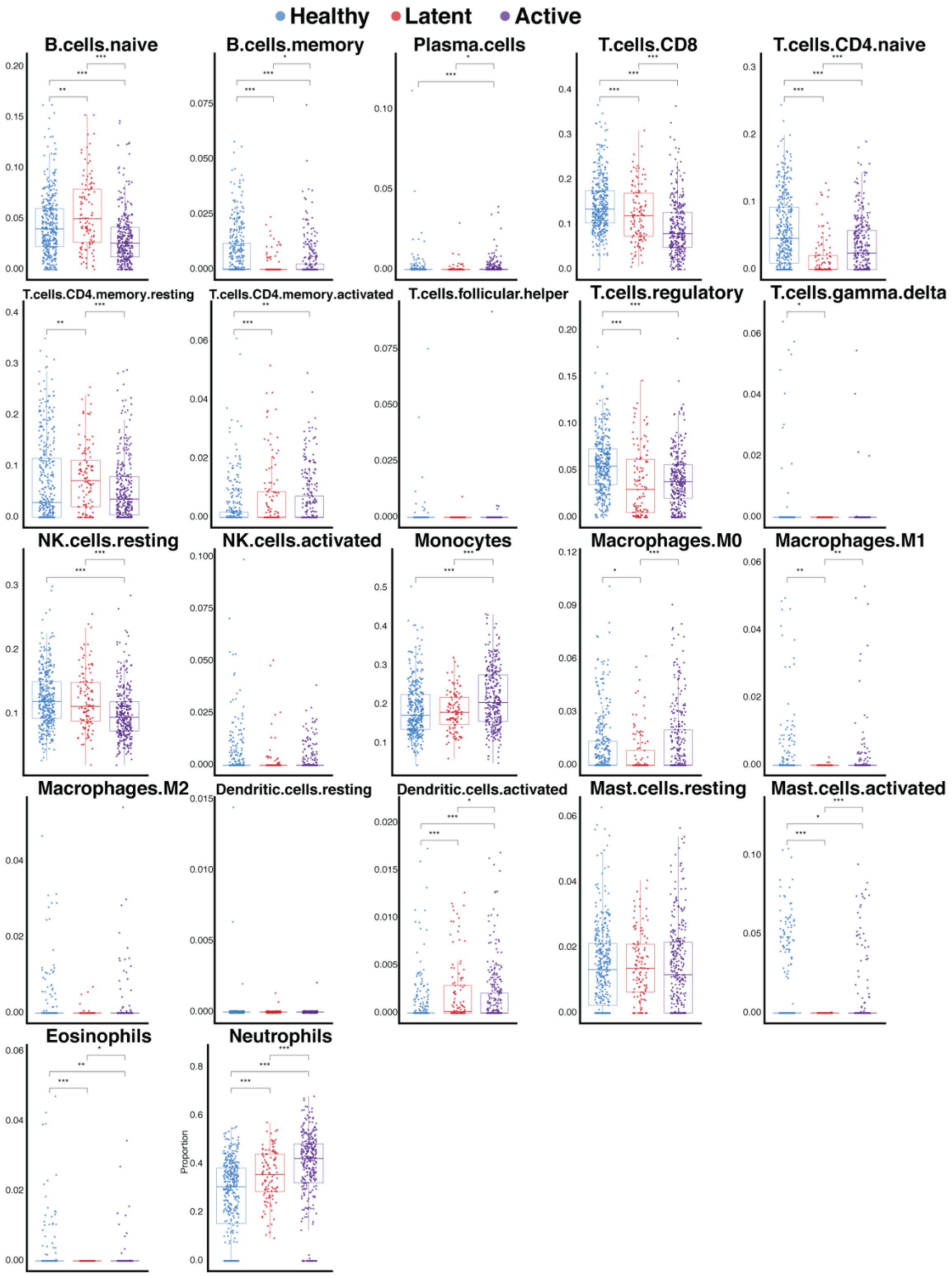


**Matrix: LM22**
**Data Type: RNA-Seq**

**Figure S5 Statistical deconvolution of 13 clinical cohorts of active and latent TB infections.** Box plots of TB infection status after deconvolution with our MCCS curated matrix for microarray (A) and RNA-Seq studies (B), the immunoStates matrix [10] for microarray (C) and RNA-Seq (D) studies and the LM22 matrix from CIBERSORT [1] for microarray (E) and RNA-Seq (F) studies. Significance determined by Kruskal-Wallis rank sum test with $p$-value < 0.05 = *, $p$-value < 0.01 = ** and $p$-value < 0.001 = ***. Sample sizes for each disease state and data type are as follows; healthy (microarray $n$ = 88, RNA-Seq $n$ = 365), latently infected (microarray $n$ = 376, RNA-Seq $n$ = 117) and active disease individuals (microarray $n$ = 547, RNA-Seq $n$ = 306).

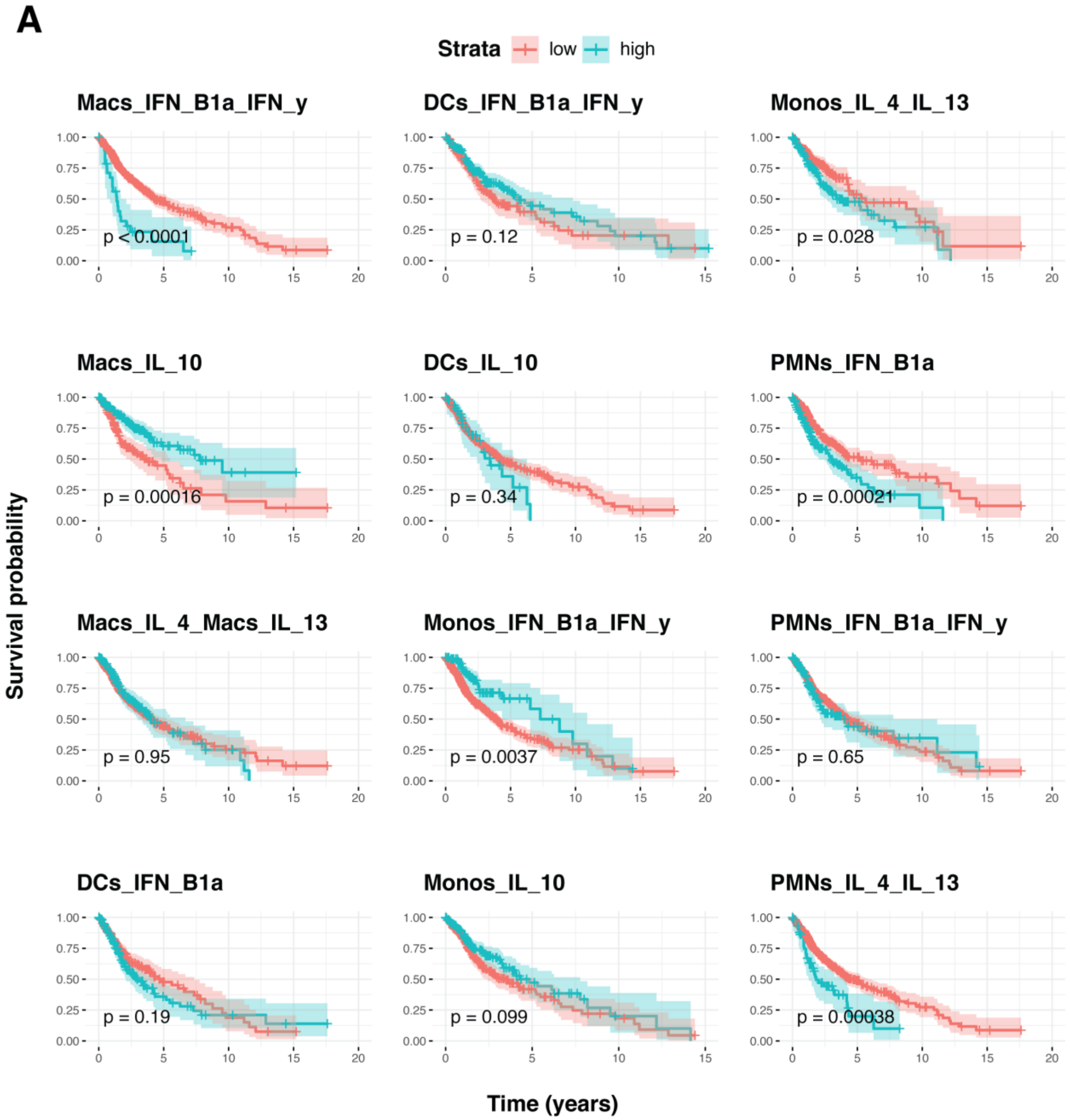


**Figure S6. MCCS signatures are linked to survival in glioma patients.** Survival analysis of 671 primary glioma tumors after cell type specific deconvolution through CIBERSORT using our MCCS signature matrix. Samples are stratified by low and high quartiles for resulting deconvolution proportion estimate. Significance determined by log rank test.

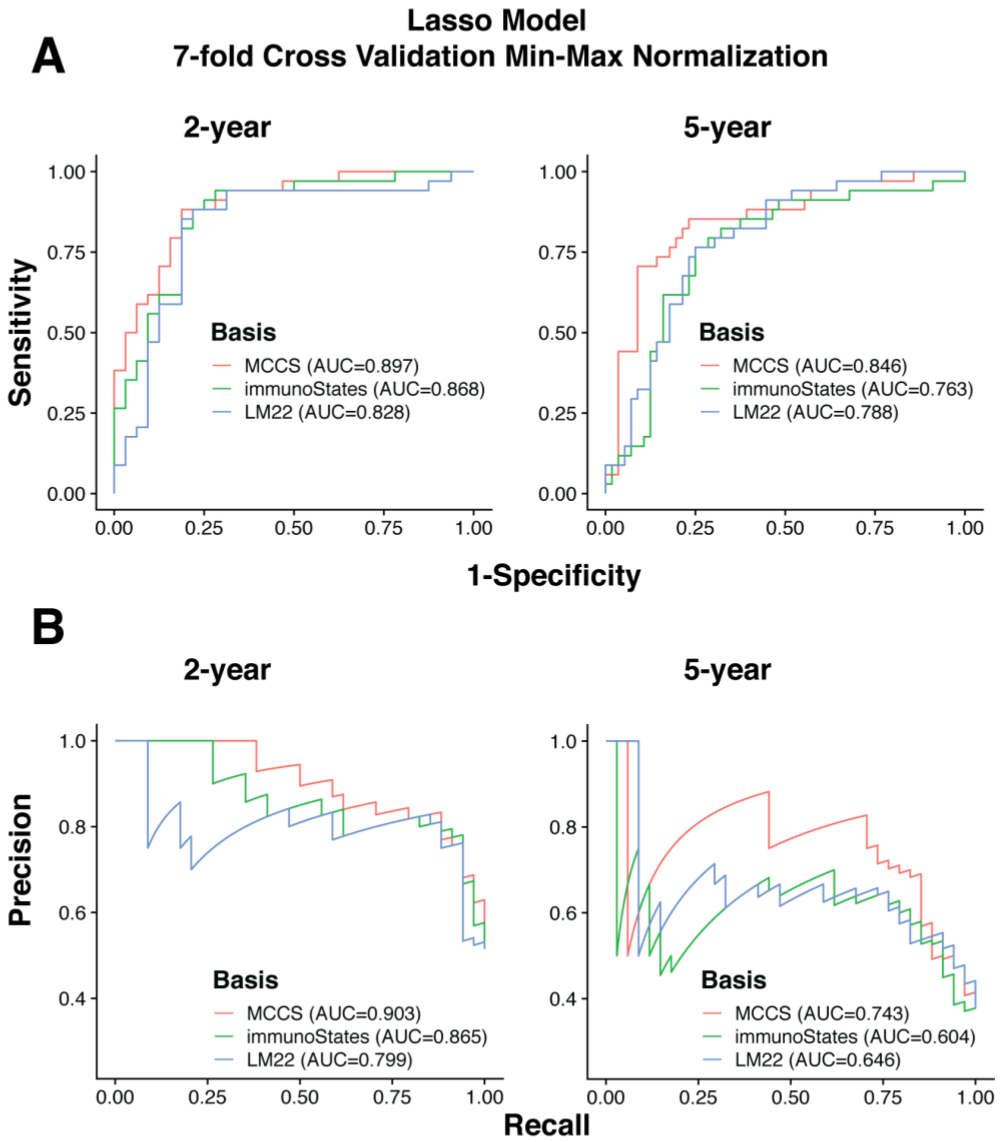


**Figure S7. MCCS gene signature is highly successful at predicting long-term survival in glioma patients.** Area under ROC curves (A) and Precision Recall curves (B) for 2 and 5-year LASSO models with 7-fold cross validation and min-max normalization. Models trained on 131 MCCS gene signature (red), 341 immunoStates signature (green) or 547 LM22 gene signature set (blue).
